# Spatiotemporal structure of intracranial electric fields induced by transcranial electric stimulation in human and nonhuman primates

**DOI:** 10.1101/053892

**Authors:** Alexander Opitz, Arnaud Falchier, Chao-Gan Yan, Erin Yeagle, Gary Linn, Pierre Megevand, Axel Thielscher, Michael P. Milham, Ashesh Mehta, Charles Schroeder

## Abstract

Transcranial electric stimulation (TES) is an emerging technique, developed to non-invasively modulate brain function. However, the spatiotemporal distribution of the intracranial electric fields induced by TES remains poorly understood. In particular, it is unclear how much current actually reaches the brain, and how it distributes across the brain. Lack of this basic information precludes a firm mechanistic understanding of TES effects. In this study we directly measure the spatial and temporal characteristics of the electric field generated by TES using stereotactic EEG (s-EEG) electrode arrays implanted in cebus monkeys and surgical epilepsy patients. We found a small frequency dependent decrease (10%) in magnitudes of TES induced potentials and negligible phase shifts over space. Electric field strengths were strongest in superficial brain regions with maximum values of about 0.5 mV/mm. Our results provide crucial information for the interpretation of human TES studies and the optimization and design of TES stimulation protocols. In addition, our findings have broad implications concerning electric field propagation in non-invasive recording techniques such as EEG/MEG.

## Introduction

Transcranial electric stimulation (TES) with weak currents is an emerging non-invasive technology for modulating brain function, with the goals of investigating causal influences in neural systems and of effecting beneficial changes in cognition and behavior^1^. Promising demonstrations of the ability to up-and down-regulate neuronal excitability^2^, ^3^, entrain spontaneous oscillatory activity^4^^−^^6^, alter cognitive performance^7^, and impact pathologic psychiatric processes^8^, are rapidly increasing momentum for TES application in the neuroscientific and clinical realms alike. In-vitro and in-vivo studies using slice cultures and rodent models^9^^−^^13^ have demonstrated the ability of weak electric fields to affect firing rates and oscillations of neural populations. However, the translation of these findings to the human brain is not straightforward, as brain folding is much more complex, and complications arise from currents passing the scalp, skull and cerebrospinal fluid (CSF) before reaching the cortex. As a result, the strength, distribution and orientation of the electric field generated by TES approaches, such as transcranial direct current stimulation (tDCS) and transcranial alternating current stimulation (tACS) remain largely unknown.

Studies focused on the interpretation of local field potentials have successfully measured the electric properties of brain tissues on a small spatial scale (i.e., in a 5mm patch)^14^. However, it is unclear how those findings generalize to electric field measurements at a full-brain level. Moreover, these studies concerned the measurement of fields generated using local intracranial current stimulation, as opposed to those caused by extracranial stimulation, leaving a wide gap in our mechanistic understanding. Some have attempted to bridge this gap by measuring the electric fields during TES using phantoms^15^, though it is not clear how faithfully an agar-filled volume can represent the CSF-filled intracranial space and the complex brain in the human. Beyond the measurement of static electric fields for tDCS, additional questions arise for tACS, where the impact of brain tissue properties on the temporal dynamics of electric fields produced by oscillating stimulation currents are incompletely understood. For example, earlier studies suggest that significant inhomogeneity of conductivity and permittivity in brain tissue gives rise to filtering of potential fields over space in a way that distorts the frequency content of the original signal and causes a phase/time shift of field potential components over space^16^, ^17^. While this conclusion has been questioned^14^, delineating any of such effects would be critical to the practice of applying tACS so that it meshes with or perturbs ongoing oscillations at specific locations in the brain. The imprecise understanding of the specific properties of electric fields generated in the brain using scalp-applied TES limits principled efforts to optimize stimulation protocols, as well as the interpretation of experimental results.

The goal of this study was to provide comprehensive measurements of the spatial and temporal variations in the properties of electric fields (i.e., strength, direction) generated using TES in a non-human primate (NHP) brain. Because our target was the intracranial electric field, rather than the brain’s response to stimulation, we opted to streamline our approach by studying the subjects under anesthesia. The NHP model is ideally suited for such measurements due to the feasibility of strategically implanting stereotactic-EEG (s-EEG) electrodes in a manner that can sample a broad range of brain areas, the ability to perform repeated measurements, and the reasonable approximation of the brain and skull structures of nonhuman primates to those of humans. Complementing these NHP measurements, we capitalized on the availability of neurosurgical patients implanted with the same type of s-EEG electrodes. While the electrode placements were solely based on clinical considerations, the recordings nonetheless allowed us to evaluate how well specific aspects of our non-human primate findings generalize to humans. Although the present study was primarily motivated by the challenges faced by TES, our findings also have broad implications for our understanding of electric field propagation within the brain, and source localization based on EEG/MEG^18^, as both face the same fundamental biophysical limits imposed by the scalp, skull and brain as volume conductors.

## Methods

### Monkey recordings

All procedures were approved by the IACUC of the Nathan Kline Institute for Psychiatric Research. Recordings were carried out in accordance with the approved guidelines. Two Cebus monkeys (one male, 13 years and 4.1 kg and one female, 11 years and 2.9 kg) were implanted with MRI-compatible (Cilux) headposts. In the first monkey three electrodes (Adtech) with a total of 32 contacts (5 mm spacing) were permanently implanted through a skull incision over the left occipital cortex. In the second monkey, the same three electrodes were similarly implanted, along with an additional 10-contact array. Recording electrodes were oriented from posterior to anterior with medial prefrontal cortex, frontal eye field and hippocampus as target regions. In the second monkey the additional electrode targeted the geniculate complex of the thalamus. Exact electrode positions were identified on a post-implantation MR image and registered to the pre-implantation image. In multiple sessions, s-EEG was recorded during TES using a Brain AMP MR amplifier (Input Impedance 10 MOhm, Common-mode rejection > 90 db, high-pass frequency 0.016 Hz/10s, low-pass frequency 250Hz, measuring range +- 16.384mV, resolution 0.5 muV/bit, Brain Products) with a sampling rate of 500 Hz. The frequency-magnitude response of the EEG system was tested using a function generator and a linear ohmic resistor to derive a correction factor for a slight dampening at higher frequencies (see Supplementary Material, Fig. 1). Ground (right temple) and reference electrodes (left temple) were attached on the scalp. For a detailed analysis regarding the effect of the position of the reference and ground electrode see Supplementary Material (Supplementary Figures 5-7). During recording sessions the two monkeys were anesthetized with ketamine 10 mg/kg, atropine 0.045 mg/kg, and diazepam 1 mg/kg followed by 2% isoflurane. Small round stimulation electrodes (3.14 cm2, Ag/AgCl with conductive gel (SigmaGel)) were used in all sessions and transcranial electrical stimulation was applied using the Starstim system (Neuroelectrics, current controlled stimulation). The position of stimulation electrodes was chosen over the left occipital cortex and middle forehead to create electric fields with a large component along the direction of the implanted electrode arrays (Fig. 3C). The current output of the stimulator was tested with a linear ohmic resistor and a minimal decrease for high frequencies was corrected for in results (see Supplementary Material, Fig. 1). To investigate the time course of electric fields during tACS and to test for possible frequency dependent effects on field strength or phase shifts, we parametrically varied the frequency of stimulation currents from 1 Hz to 150 Hz. We tested 21 frequencies (1-10 Hz in 1Hz steps, 10Hz – 100 Hz in 10 Hz steps and 125 + 150 Hz). The frequencies were tested with ca. 30 - 60s rest time between each frequency measurement. Frequencies were applied in randomized order as stimulation itself can change the impedance of electrodes/skin over time^19^, thus performing measurements with increasing/decreasing frequencies could lead to systematic errors. The impedance of the stimulation electrodes was monitored after every five frequencies measured to ensure that no large changes occurred over the course of the experiment. Electrode impedance was below 5 kOhm during the recordings. In one session we recorded from two additional scalp needle electrodes attached on the left and right side along the head midline to test whether phase shifts might occur through the current passing the skull. We applied alternating currents at 200 muA in the 1^st^ monkey and 100 muA in the 2^nd^ monkey (which had thinner temporalis muscles and a smaller head size) to ensure a similar voltage range, through two electrodes (left occipital cortex, middle forehead) for 30s for at each frequency. Intensities were chosen to achieve a high signal to noise ratio while keeping within the dynamic range of the amplifier, as well as to reduce strain on the monkeys for extended measurement sessions. Each measurement series was repeated once immediately following the first series.

**Figure 1:**
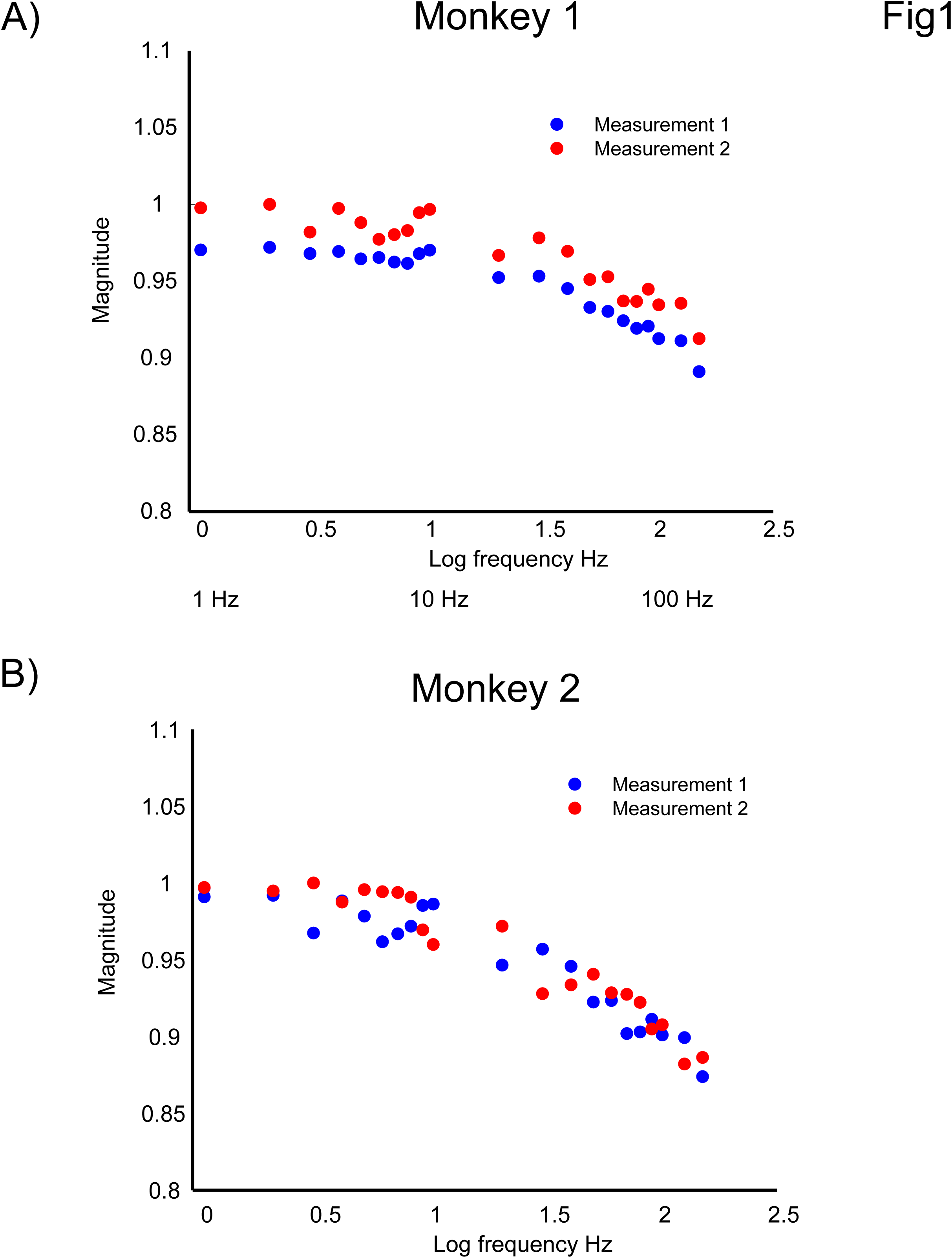
Bode plot illustrating the frequency dependency of the magnitude of TES induced electric potentials measured in Monkey 1 **A)** and Monkey 2 **B)**. All shown results are corrected for the dampening of the stimulation and recording system. Normalized mean magnitude over all contacts in dependence of stimulation frequency (log10 units) from 1Hz – 150 Hz for two repeated measurements. A slight decrease in magnitude of up to 10% is visible for higher stimulation strengths.

**Figure 3:**
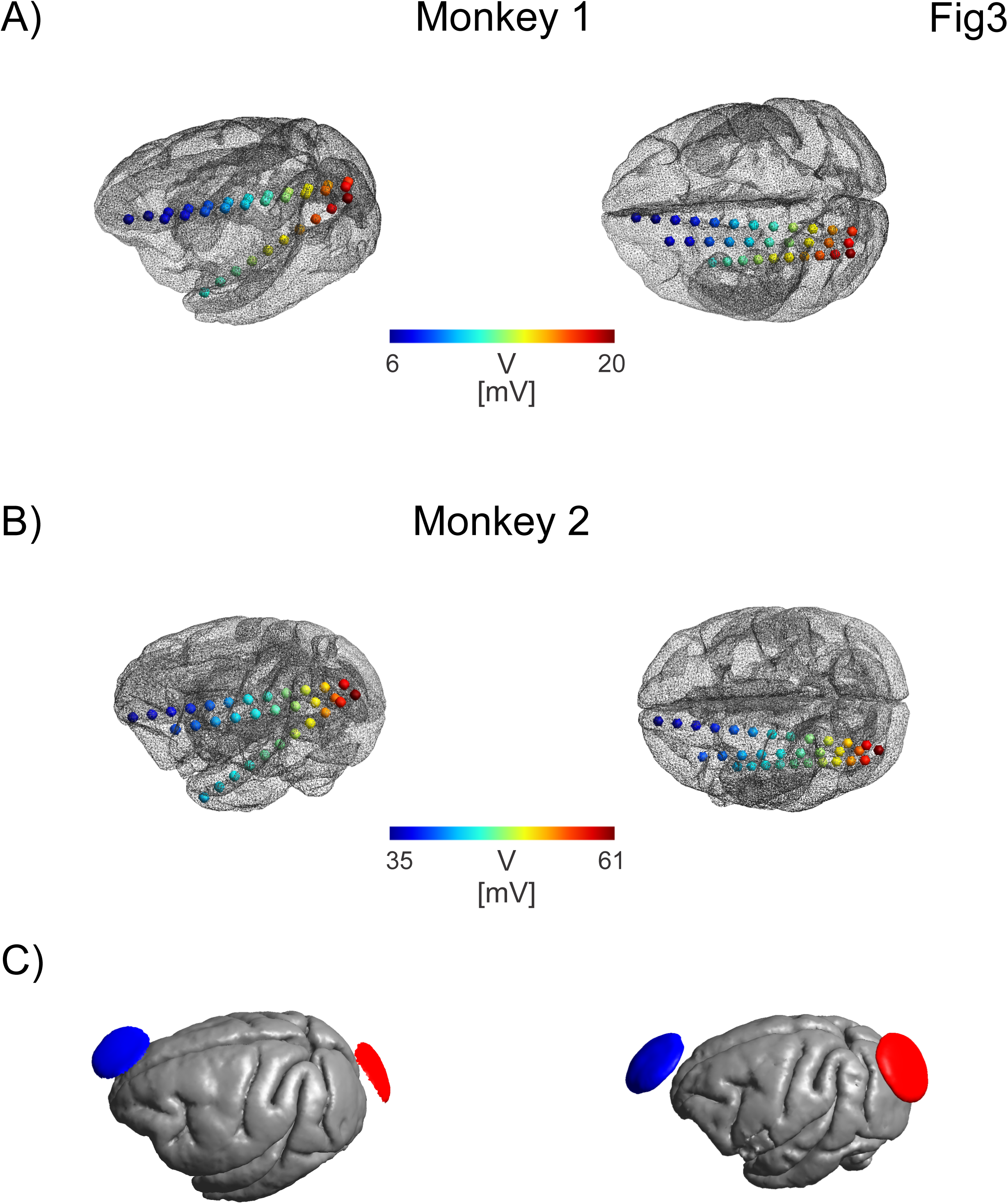
Intracranial potential distribution for monkey 1 **A)** and monkey 2 **B)**. Shown is the measured electric potential (in mV scaled for a stimulation intensity of 1mA, measured at 1Hz) at different electrode contacts implanted in the left hemisphere. Stimulation electrodes were attached over the left occipital cortex and middle forehead and their locations are indicated with red and blue arrows for both monkeys. A continuously changing posterior - anterior gradient in the electric potential is visible. **C)** Stimulation electrodes displayed over the cortical surface for Monkey 1 (left) and Monkey 2 (right).

### Patient recordings

Two patients (one female, one male, both right-handed, ages 35 and 29, respectively) with refractory epilepsy participated in the study while undergoing presurgical monitoring at North Shore University Hospital. Seizure onset zones were multifocal for patient 1 (right hippocampal/temporal, left temporal, left orbitofrontal) and left amygdala and hippocampus for patient 2. All experimental protocols were approved by the Institutional Review Board of the North Shore University Hospital. All recordings were carried out in accordance with the approved guidelines. Both patients provided informed consent as monitored by the local Institutional Review Board and in accordance with the ethical standards of the Declaration of Helsinki. One patient was implanted with bilateral s-EEG electrodes (Adtech Medical Instrument Corp.) and one patient with left subdural grid, strip, and depth electrodes (Integra Lifesciences Corp.), with the number and placement of electrodes determined solely by clinical requirements. The reference electrode was placed at midline between skull and scalp for both patients. Electrode positions were identified on post-implantation CT scan^20^ and registered in a two-step procedure to the post-implantation MR and then to the pre-implantation MR^21^. Patients were monitored until sufficient data were collected to identify the seizure focus, 8-12 days. Continuous intracranial video-EEG monitoring was performed with standard recording systems (XLTEK EMU 128 LTM System) with a sampling rate of 500 Hz. In a single session patients were stimulated with TES. Two saline-soaked sponge electrodes (25cm^2^) were attached to the scalp over the left and right temple (bilateral montage) and a 1Hz alternating current of 1mA was applied in one run (using Starstim) for 2 min with a ramp up/down of 10 s. Electrode locations were chosen in close proximity to areas with good coverage by recording electrodes. Electrode impedance was kept below 10 kOhm during the recordings. Patients were instructed to rest during tACS application. As would be expected, patients reported mild skin sensations during the application of tACS.

### Data analysis

Data analysis was identical for patient and monkey recordings. First, we subtracted from each channel its mean voltage over a time interval of 1s right before stimulation onset to correct for possible baseline differences between channels unrelated to stimulation. An example of the baseline correction is shown in Supplementary Figure 9. To investigate possible frequency dependent effects of the stimulation current on magnitude and phase of the intracranial currents, we computed a Bode plot which explores the frequency response of the system. For that we computed the Fast Fourier Transform (FFT) using 16.4s of data (2^13^ data points) for each channel. An illustration of the Fourier analysis can be found in Supplementary Figure 2. To minimize FFT scalloping loss, which can exert a frequency dependent effect on estimated FFT magnitude, we applied a flat-top window function to the data^22^. We extracted the magnitude and phase for the maximum frequency (which was identical with the stimulation frequency for each frequency tested). Magnitude was defined as the absolute value of the complex fourier value at the peak frequency. Average magnitudes were computed over all channels (scalp electrodes were analyzed separately) for each frequency and normalized to the maximum amplitude. In the following, we computed phase differences for the peak frequency between all possible channel combinations. We excluded phase differences from channels with very small FFT magnitudes relative to the maximally observed amplitudes as in those channels the phase estimation was not reliable. Then we computed the phase differences modulo π (180 degree) and centered the distribution between - π/2 and + π/2. We did this because depending on the placement of the reference electrode, a phase reversal of π could occur at some channels caused by negligible phase differences and the periodicity of the phase function (see Supplementary Material, Figure 5). Those 180-degree phase reversals arise due to the position of the reference electrode and not due to capacitive effects of the tissue (see Supplementary Material, Figures 5-7). From the phase difference histograms we then calculated the mean values of absolute phase differences.

In a second analysis we estimated the electric field strength during TES. For that we calculated the numerical gradient using the symmetric difference quotient (using a simple two point estimation leads to similar results) of the potential at its peak up-phase along the contacts for each implanted electrode. For the patient with grid electrodes, we computed the gradient along both grid axes and combined them using vector addition. All values for the potential and electric field strength were scaled to a stimulation current of 1mA and mean and standard error of the mean were computed over five stimulation cycles. Note that with this method we can only estimate the electric field component along the measurement (electrode) vectors (projection of the electric field). While we tried to use montages that resulted in electric fields with a large component along the electrode directions, the strength of the “true” electric field is likely somewhat larger than the measured field. In addition to field strength, we estimated the spatial extent of intracranial electric fields. First, we identified those electrode contacts that exhibited electric field strengths larger than 50% or 25% of the maximum recorded field strength; this was carried out separately for each implanted linear electrode array or ECoG grid. Then, for each electrode array, we computed the maximum distance between those contacts that met the cutoff criteria.

Based on the pre-implantation MR images, we reconstructed the cortical surfaces to visualize the experimental recordings in a 3D model for both monkeys and patients.

## Results

The analysis of the frequency response function indicated that the mean potential magnitude decreases as a function of stimulation frequency (Fig. 1), with a maximum decrease of around 10% for the highest stimulation frequency tested (150 Hz). While magnitudes were largely stable for slow frequencies up to 15 Hz, there was a continuous decline from 15 to 150 Hz. Negative correlations of *r*(19) = −0.98, *p* < .001 and *r*(19) = −0.94, *p* < .001 between frequency and magnitude were found for monkeys 1 and 2, respectively. This frequency dependent decrease of magnitude is in line with a frequency dependent increase of conductivities of the head tissue or electrodes, as a smaller voltage is needed to pass the same amount of current in a better conducting medium. We observed only very small phase differences, up to a few degrees, between electrodes (Fig. 2), indicating that capacitive effects that would produce phase differences are quite small. Results were consistent between monkeys. Between frequency and phase differences positive correlations of *r*(19) = 0.87, *p* < .001 and *r*(19) = 0.98, *p* < .001 were found for monkeys 1 and 2. Phase differences in the patients (not shown) were generally in the same small range as in the monkeys. For the scalp electrodes we observed the same mild frequency dependence of the FFT magnitude, and again, we observed only small phase shifts from the scalp electrodes to the intracranial electrodes. The lack of a phase difference between scalp and intracortical measurements suggests that the passage of current through the skull does not introduce appreciable phase shifts.

**Figure 2:**
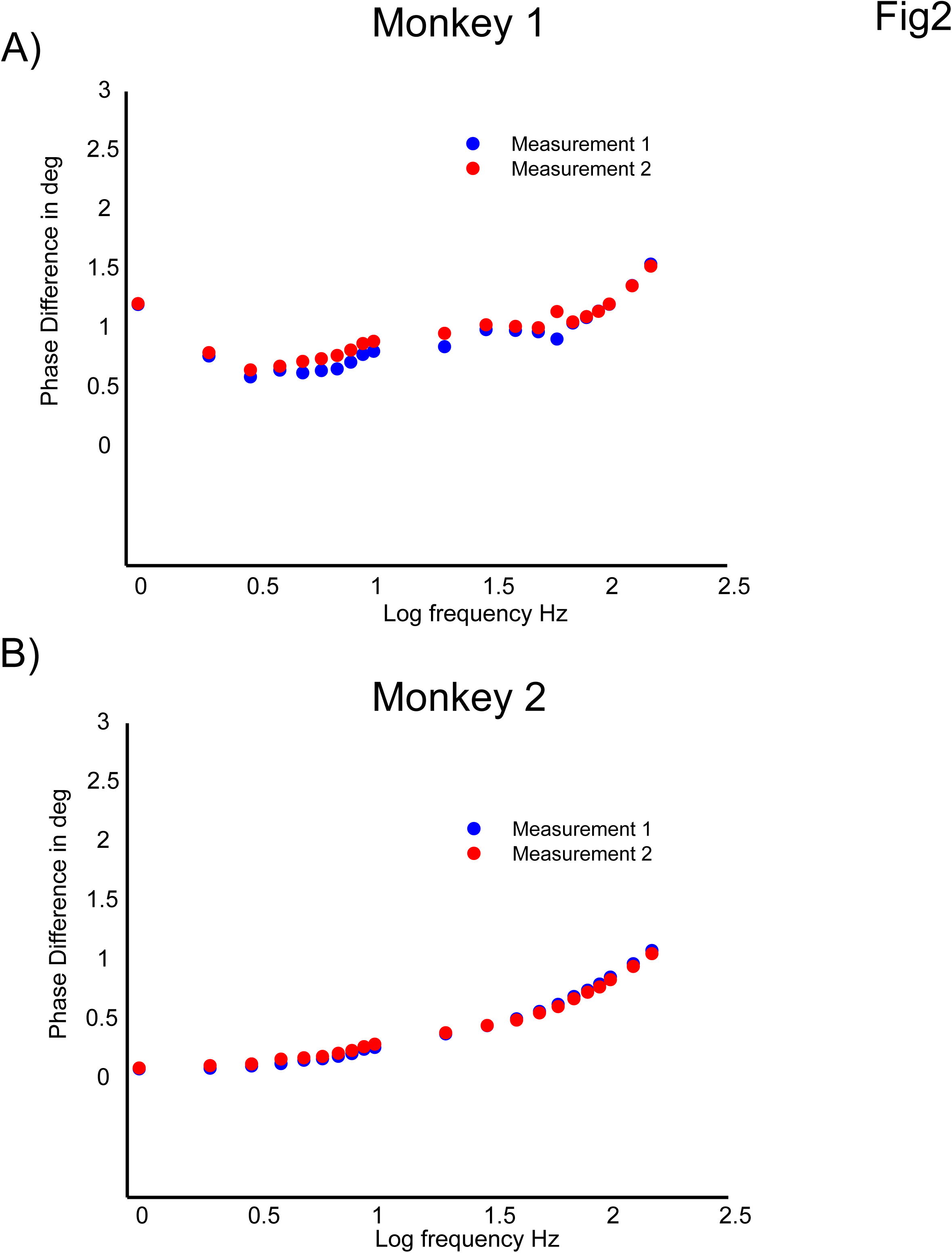
Bode plot illustrating the frequency dependency of phase differences of TES induced electric potentials measured in Monkey 1 **A)** and Monkey 2 **B)**. Mean phase differences (degree) between all combinations of electrode contacts are shown in dependence of stimulation frequency (log10 units) from 1Hz – 150 Hz for two repeated measurements. Weak phase differences around 1-2 degrees were observed for both monkeys.

S-EEG recordings revealed potential distributions with continuously varying gradients between the stimulation electrodes both in monkeys and humans (Figs. 3 + 4), however, subdural grid array recordings measuring the potential distribution at the outer brain surface (Fig. 4b) yielded a more complex gradient. The rapid change of electric field strength over space in Patient 2 is likely related to a quickly changing radial (inwards) to tangential (along the cortical surface) electric field component. Our electric field measurements are only sensitive in the tangential component, resulting in the pattern observed here. Regarding electric fields, largest magnitudes were generally found in superficial sites near the stimulation electrodes but not necessarily confined to the outermost contacts (Figs. 5 + 6).

**Figure 4:**
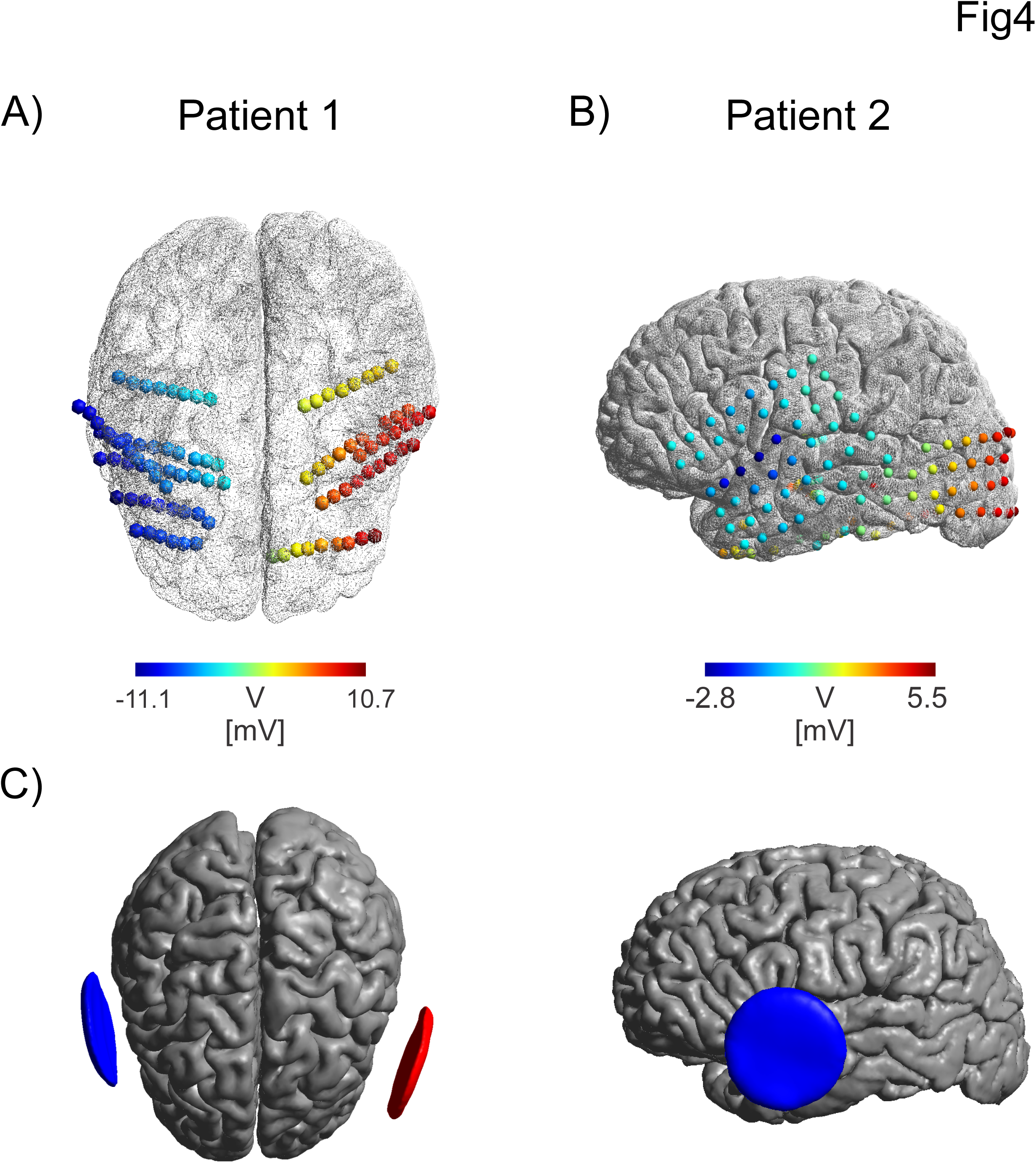
Intracranial potential distribution for Patient 1 **A)** and Patient 2 **B)**. Measured electric potential (in mV scaled for a stimulation intensity of 1mA) at different bi-hemispheric stereotactic EEG electrode contacts (Patient 1) or surface ECoG grid on the left hemisphere (Patient 2). Stimulation electrodes were attached bilaterally over the left and right temple in both patients and indicated with red and blue arrows. A continuously changing left - right gradient in the electric potential is visible for Patient 1. For Patient 2 a sharp change in potential is found close to the left stimulation electrode. Continuously increasing potentials are found with increasing distance to the stimulation electrode. Note the large potentials found in the occipital region are due to the lack of electrode coverage on the right hemisphere which would exhibit even higher values. **C)** Stimulation electrodes shown over the cortical surface for Patient 1 (left) and Patient 2 (right, other cross hemispheric electrode not visible).

**Figure 5:**
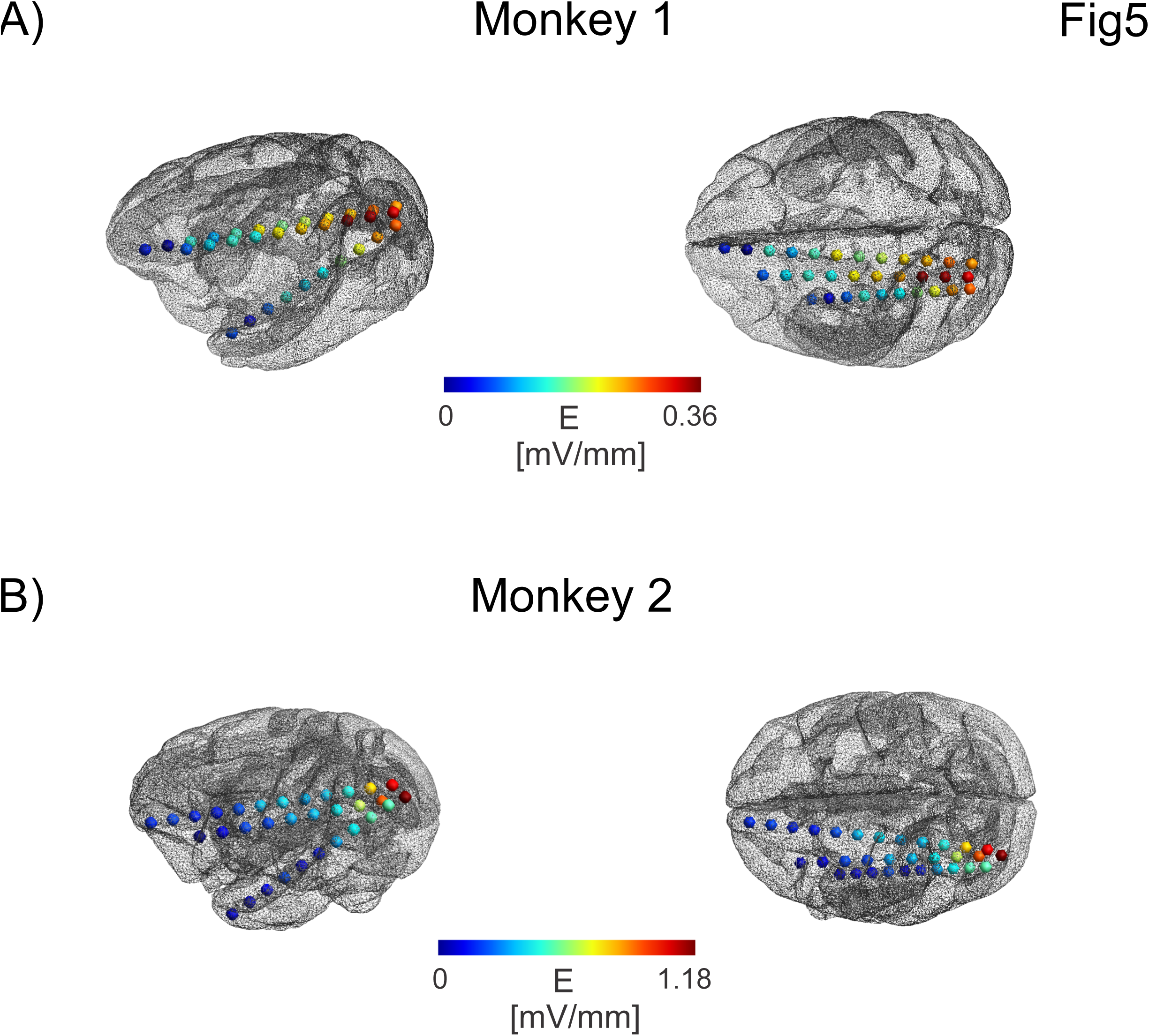
Intracranial electric field distribution for monkey 1 **A)** and monkey 2 **B)**. The position of the stimulation electrodes on the scalp are indicated with red and blue arrows. The electric field projection along the electrodes (in mV/mm scaled for a stimulation intensity of 1mA) shows an intricate pattern with high electric fields close to the occipital stimulation electrode (monkey 1). Note that the strongest electric field strength was found not at the most superficial recording electrode but at an electrode a bit deeper in the cortex. The weak electric field strengths near the frontal stimulation electrode are likely due to the larger distance to the frontal electrode. For the second monkey strongly enhanced electric field strength occurred at one electrode. Possible reasons are smaller head size and reduction in muscle tissue that can lead to larger field strengths. Also the contact with highest electric field strength was outside the brain, possibly explaining the large field strength.

**Figure 6:**
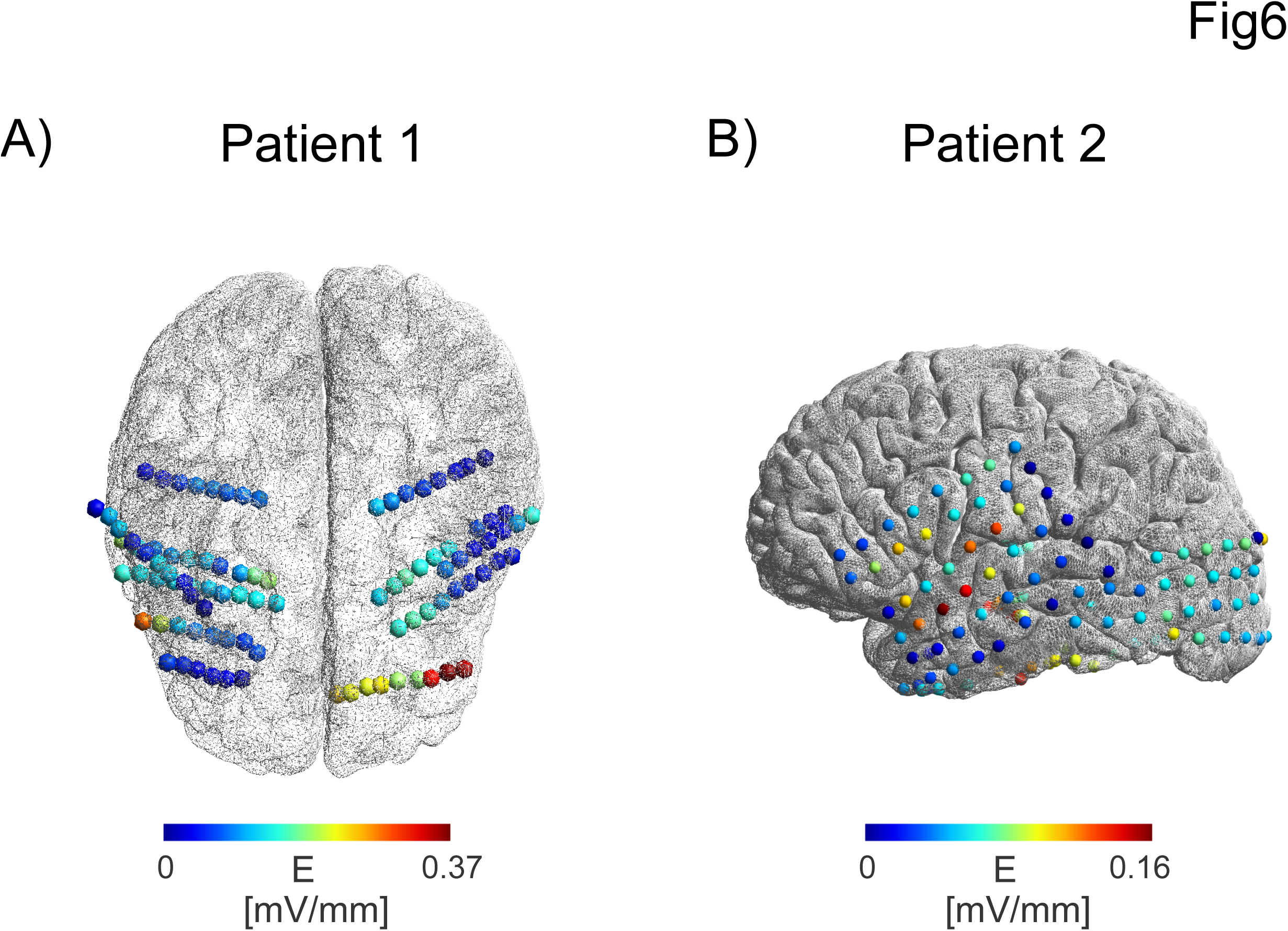
Intracranial electric field distribution for Patient 1 **A)** and Patient 2 **B)** at different bihemispheric stereotactic EEG electrode contacts (Patient 1) or surface ECoG grid on the left hemisphere (Patient 2). The position of the stimulation electrodes on the scalp are indicated with red and blue arrows. Shown is the electric field projection in mV/mm scaled for a stimulation intensity of 1mA. Highest electric field strength was found at contacts close to the stimulation electrodes (bilaterally) with decreasing strength for increasing depth in the brain for Patient 1. In Patient 2 highest electric field strength was found near the contacts close to the stimulation electrode (left hemisphere) and decreasing values at more remote electrodes.

Maximum electric strengths were found to be 0.358 +/− 0.001 mV/mm for the first monkey (median electric field 0.21 mV/mm) and 1.173 +/− 0.003 mV/mm for the second monkey (median electric field 0.39 mV/mm). The first monkey was a male with a much larger muscle mass overlaying the skull than the second, a female monkey, highlighting the fact that individual anatomical factors result in different electric field strengths. Maximum electric field strengths for the patients were found to be 0.360 +/− 0.008 mV/mm (Patient 1) (median electric field 0.098 mV/mm), 0.163 +/- 0.007 mV/mm (Patient 2) (median electric field 0.059 mV/mm).

We evaluated the spatial extent of electric fields (larger than 50% or 25% of the maximum electric field) as follows: Monkey 1: 90 mm (50% of Max) and 110 mm (25% of Max), Monkey 2: 20 mm and 60 mm. Patient 1: 39.6 mm and 45.7 mm, Patient 2: 47 mm and 74.6 mm. The differences in spatial extent observed across cases were predictable given that the electrodes were not placed in the same locations or orientations. However, the findings for the monkey are more notable, as their placement was relatively similar in the two subjects. This again suggests that other factors, such head size, muscle tissue thickness or skull integrity may contribute to inter-individual differences in field properties.

## Discussion

Comprehensive evaluation of the intracranial electric field during TES in nonhuman primates and human neurosurgical patients provided empirical support for common assumptions about the electric properties of the head tissue^23^, ^24^ and provided novel insights with implications that extend beyond TES. More specifically, we confirmed that the electric fields generated during TES do behave in a linear ohmic manner, with small capacitive components on a mesoscopic or macroscopic scale, alleviating several concerns that could greatly complicate dosing. We observed a frequency dependent attenuation of the voltages generated, up to 10%, for tACS frequencies exceeding 15Hz. This phenomenon can be explained by frequency dependent increases in conductivity^25^ (i.e., for a current controlled stimulation a smaller voltage is needed to achieve a fixed current strength in a higher conducting medium). Thus, while likely not critical, studies comparing physiological and behavioral responses to different stimulation frequencies could account for the differences in electric field strength arising from specific stimulation frequencies, especially when considering higher frequencies. Capacitive effects that could result in phase shifts were generally small. This finding is crucial for tACS stimulation protocols that aim to exploit the phase relationship between injected currents and ongoing rhythmic brain activity^26^^−^^28^. While previous publications reported larger capacitive effects of brain tissue e.g. ^29^^−^^31^ compared to this study, our measurements were conducted in-vivo with a “4-electrode setup” (two separate stimulation and recording electrodes) minimizing capacitive effects at the electrode-electrolyte interface of the recording electrodes. In line with our results, small phase differences were also reported in Logothetis, Kayser and Oeltermann ^14^ using a similar measurement setup on a smaller spatial scale. We did not measure the phase differences between tACS stimulation currents and recorded voltages, but only between voltages at different contacts. For the stimulation electrode on the scalp, phase differences could occur at the electrode-electrolyte interface. Thus our results are primarily applicable to the volume conduction problem of currents passing through scalp, skull and brain. While we did not observe large capacitive effects on the mesoscopic scale, they could possibly occur on a microscopic scale e.g. due to membrane capacitance of neurons. Future work, using the appropriate apparatus for recordings cellular level phenomena would be required to evaluate potential effects at the microscale. Interestingly, however, phase shifts were not observed even for close-by electrode pairs located in homogeneous parts of brain white matter. This indicates that membrane capacitances do not result in strong deviations from a purely ohmic behavior on the spatial scale which is relevant for shaping the gross field distribution. Obviously, our conclusions only hold for the measured frequency range, which was restricted by the technical parameters of the stimulation and recording equipment. However, it is noteworthy that prior measurements often reported the strongest deviations from an ohmic behavior in the frequency range tested here^25^, ^31^, ^32^, highlighting the importance of the data reported. One difference between our measurement and others performed previously (e.g. Gabriel, Peyman and Grant ^32^) is that no acute strain or compression of the tissue is induced by the recording setup we used.

Our measurements also have direct implications for interpretation of the EEG signal based on the reciprocity theorem^33^, ^34^. Briefly, this theorem states that the electric field at a certain location in the brain arising from an imposed current through two scalp electrodes is equivalent to the voltage measured through the same two scalp electrodes resulting from a dipole at the same brain location. Based on our measurements, we conclude that a dipole in the brain for high frequencies will result in lower measured voltages at the scalp electrodes. Nevertheless the moderate size of this lowpass filtering effect cannot alone explain the 1/f behavior^35^ observed in recordings of brain oscillations.

Our results show that maximum electric fields in humans reach up to 0.5 mV/mm for 1mA stimulation currents which is in the range predicted by modeling studies^36^, ^37^. This is a lower bound for which actual neural entrainment has been found in in-vitro studies^13^, ^38^ and below the threshold of 1mV/mm in rodent studies^4^. It is thus not completely clear how mechanisms derived from in-vitro and animal studies can be applied to human studies due to the limited field strength induced in human brains. Future efforts could be made to induce higher electric field strengths in a non-painful and safe manner; e.g., using kHz currents as a carrier frequency for low frequency oscillations. Input currents used in human studies could be matched to achieve electric field strengths shown to be effective in in-vitro settings. We found substantial inter-individual differences in electric field strength which might underlie some of the variability in response between participants to TES^39^, ^40^ and support the suggestion that dosing being individualized. The spatial extent of the electric field was estimated to range over several cm in both monkeys and patients. This is in line with predictions from modeling studies showing electric fields extend across several gyri^37^. Novel montages using multiple stimulation electrodes have the potential to increase spatial specificity of delivered electric fields and could be studied in future research^41^. While due to technical limitations we could not directly measure DC currents, we expect that the findings at the low frequency limit for our measurements apply for tDCS as well, especially under the aspect of small frequency dependent effects. Future studies should test how well observed electric field distributions accord with predictions of computational models^42^. While a comprehensive evaluation and fitting of finite element models to the measured data is outside the scope of this study, such an approach seems promising especially with respect to optimizing montages and stimulation protocols to produce more focal fields^41^. In addition, our results are also crucial for the precise tuning of closed loop systems that can tailor the stimulation current and phase to ongoing oscillations^43^. To target ongoing oscillations in a phase-specific manner, changes in phase or magnitude that occur when currents pass through the skull and brain tissues need to be taken into account.

**Table 1.**
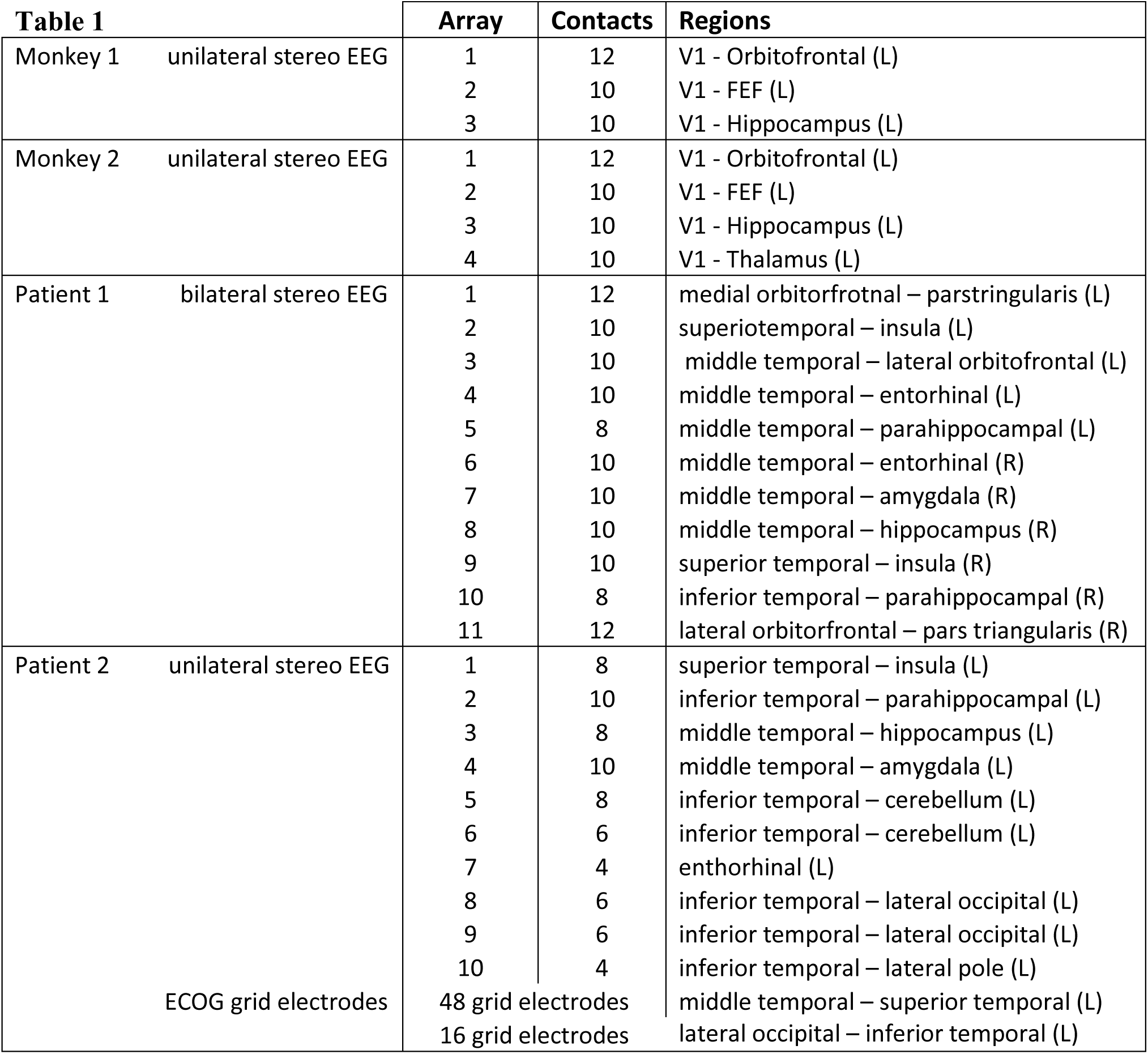
Overview of implanted electrodes (number and type) and covered brain regions.

## Author Contributions Statement

“A.O, A.F, CG.Y, E.Y, G.L, P.M, A.M, C.S performed experiments. A.O analyzed data and prepared all figures. A.O, A.T, MP.M, C.S. interpreted results. A.O, A.T, M.P.M, C.S. wrote the main manuscript text. All authors reviewed the manuscript.”

## Additional Information

## References

1 Paulus, W. Transcranial electrical stimulation (tES - tDCS; tRNS, tACS) methods. Neuropsychological rehabilitation 21, 602–617 (2011).

2 Nitsche, M.A. & Paulus, W. Excitability changes induced in the human motor cortex by weak transcranial direct current stimulation. The Journal of physiology 527 Pt 3, 633–639 (2000).

3 Nitsche, M.A. & Paulus, W. Sustained excitability elevations induced by transcranial DC motor cortex stimulation in humans. Neurology 57, 1899–1901 (2001).

4 Ozen, S., et al. Transcranial electric stimulation entrains cortical neuronal populations in rats. The Journal of neuroscience: the official journal of the Society for Neuroscience 30, 11476–11485 (2010).

5 Ali, M.M., Sellers, K.K. & Frohlich, F. Transcranial alternating current stimulation modulates large-scale cortical network activity by network resonance. The Journal of neuroscience: the official journal of the Society for Neuroscience 33, 11262–11275 (2013).

6 Helfrich, R.F., et al. Entrainment of brain oscillations by transcranial alternating current stimulation. Curr Biol 24, 333–339 (2014).

7 Coffman, B.A., Clark, V.P. & Parasuraman, R. Battery powered thought: enhancement of attention, learning, and memory in healthy adults using transcranial direct current stimulation. NeuroImage 85 Pt 3, 895–908 (2014).

8 Kuo, M.-F., Paulus, W. & Nitsche, M.A. Therapeutic effects of non-invasive brain stimulation with direct currents (tDCS) in neuropsychiatric diseases. NeuroImage 85, Part 3, 948–960 (2014).

9 Bindman, L.J., Lippold, O.C. & Redfearn, J.W. Long-lasting changes in the level of the electrical activity of the cerebral cortex produced bypolarizing currents. Nature 196, 584–585 (1962).

10 Bikson, M., et al. Effects of uniform extracellular DC electric fields on excitability in rat hippocampal slices in vitro. The Journal of physiology 557, 175–190 (2004).

11 Reato, D., Rahman, A., Bikson, M. & Parra, L.C. Low-intensity electrical stimulation affects network dynamics by modulating population rate and spike timing. The Journal of neuroscience: the official journal of the Society for Neuroscience 30, 15067–15079 (2010).

12 Frohlich, F. & McCormick, D.A. Endogenous electric fields may guide neocortical network activity. Neuron 67, 129–143 (2010).

13 Anastassiou, C.A., Perin, R., Markram, H. & Koch, C. Ephaptic coupling of cortical neurons. Nature neuroscience 14, 217–223 (2011).

14 Logothetis, N.K., Kayser, C. & Oeltermann, A. In vivo measurement of cortical impedance spectrum in monkeys: implications for signal propagation. Neuron 55, 809–823 (2007).

15 Kim, D., et al. Validation of Computational Studies for Electrical Brain Stimulation With Phantom Head Experiments. Brain stimulation 8, 914–925 (2015).

16 Bedard, C., Kroger, H. & Destexhe, A. Modeling extracellular field potentials and the frequency-filtering properties of extracellular space. Biophysical journal 86, 1829–1842 (2004).

17 Bedard, C., Rodrigues, S., Roy, N., Contreras, D. & Destexhe, A. Evidence for frequency-dependent extracellular impedance from the transfer function between extracellular and intracellular potentials: intracellular-LFP transfer function. J Comput Neurosci 29, 389–403 (2010).

18 Michel, C.M., et al. EEG source imaging. Clinical neurophysiology: official journal of the International Federation of Clinical Neurophysiology 115, 2195–2222 (2004).

19 Hahn, C., et al. Methods for extra-low voltage transcranial direct current stimulation: current and time dependent impedance decreases. Clinical neurophysiology: official journal of the International Federation of Clinical Neurophysiology 124, 551–556 (2013).

20 Dykstra, A.R., et al. Individualized localization and cortical surface-based registration of intracranial electrodes. NeuroImage 59, 3563–3570 (2012).

21 Groppe, D.M., et al. Dominant frequencies of resting human brain activity as measured by the electrocorticogram. NeuroImage 79, 223–233 (2013).

22 Lyons, R. Reducing FFT Scalloping Loss Errors Without Multiplication [DSP Tips and Tricks]. Signal Processing Magazine, IEEE 28, 112–116 (2011).

23 Plonsey, R. & Heppner, D.B. Considerations of quasi-stationarity in electrophysiological systems. The Bulletin of mathematical biophysics 29, 657–664 (1967).

24 Miranda, P.C. Physics of effects of transcranial brain stimulation. Handbook of clinical neurology 116, 353–366 (2013).

25 Gabriel, C., Gabriel, S. & Corthout, E. The dielectric properties of biological tissues: I. Literature survey. Physics in medicine and biology 41, 2231–2249 (1996).

26 Struber, D., Rach, S., Trautmann-Lengsfeld, S.A., Engel, A.K. & Herrmann, C.S. Antiphasic 40 Hz oscillatory current stimulation affects bistable motion perception. Brain Topogr 27, 158–171 (2014).

27 Polania, R., Moisa, M., Opitz, A., Grueschow, M. & Ruff, C.C. The precision of value-based choices depends causally on fronto-parietal phase coupling. Nature communications 6, 8090 (2015).

28 Vossen, A., Gross, J. & Thut, G. Alpha Power Increase After Transcranial Alternating Current Stimulation at Alpha Frequency (alpha-tACS) Reflects Plastic Changes Rather Than Entrainment. Brain stimulation 8, 499–508 (2015).

29 Gabriel, S., Lau, R.W. & Gabriel, C. The dielectric properties of biological tissues: II. Measurements in the frequency range 10 Hz to 20 GHz. Physics in medicine and biology 41, 2251–2269 (1996).

30 Wagner, T., Valero-Cabre, A. & Pascual-Leone, A. Noninvasive Human Brain Stimulation. Annu Rev Biomed Eng 9, 527–565 (2007).

31 Wagner, T., et al. Impact of brain tissue filtering on neurostimulation fields: a modeling study. NeuroImage 85 Pt 3, 1048–1057 (2014).

32 Gabriel, C., Peyman, A. & Grant, E.H. Electrical conductivity of tissue at frequencies below 1 MHz. Physics in medicine and biology 54, 4863–4878 (2009).

33 Plonsey, R. RECIPROCITY APPLIED TO VOLUME CONDUCTORS AND THE ECG. IEEE Trans Biomed Eng 10, 9–12 (1963).

34 Rush, S. & Driscoll, D.A. EEG Electrode Sensitivity-An Application of Reciprocity. Biomedical Engineering, IEEE Transactions on >BME-16, 15–22 (1969).

35 Pritchard, W.S. The brain in fractal time: 1/f-like power spectrum scaling of the human electroencephalogram. The International journal of neuroscience 66, 119–129 (1992).

36 Datta, A., et al. Gyri-precise head model of transcranial direct current stimulation: improved spatial focality using a ring electrode versus conventional rectangular pad. Brain stimulation 2, 201–207, 207 e201 (2009).

37 Miranda, P.C., Mekonnen, A., Salvador, R. & Ruffini, G. The electric field in the cortex during transcranial current stimulation. NeuroImage 70, 48–58 (2013).

38 Radman, T., Su, Y., An, J.H., Parra, L.C. & Bikson, M. Spike timing amplifies the effect of electric fields on neurons: implications for endogenous field effects. The Journal of neuroscience: the official journal of the Society for Neuroscience 27, 3030–3036 (2007).

39 López-Alonso, V., Cheeran, B., Río-Rodríguez, D. & Fernández-del-Olmo, M. Inter-individual Variability in Response to Non-invasive Brain Stimulation Paradigms. BRAIN STIMULATION: Basic, Translational, and Clinical Research in Neuromodulation (2014).

40 Wiethoff, S., Hamada, M. & Rothwell, J.C. Variability in response to transcranial direct current stimulation of the motor cortex. BRAIN STIMULATION: Basic, Translational, and Clinical Research in Neuromodulation (2014).

41 Dmochowski, J.P., Datta, A., Bikson, M., Su, Y. & Parra, L.C. Optimized multi-electrode stimulation increases focality and intensity at target. Journal of neural engineering 8, 046011 (2011).

42 Opitz, A., Paulus, W., Will, S., Antunes, A. & Thielscher, A. Determinants of the electric field during transcranial direct current stimulation. NeuroImage 109C, 140–150 (2015).

43 Berenyi, A., Belluscio, M., Mao, D. & Buzsaki, G. Closed-loop control of epilepsy by transcranial electrical stimulation. Science 337, 735–737 (2012).

